# PALB2 maintains redox and mitochondrial homeostasis in the brain and cooperates with ATG7 to suppress p53 dependent neurodegeneration

**DOI:** 10.1101/2021.06.07.447248

**Authors:** Yanying Huo, Akshada Sawant, Yongmei Tan, Amar H Mahdi, Tao Li, Hui Ma, Vrushank Bhatt, Run Yan, Jake Coleman, Cheryl F. Dreyfus, Jessie Yanxiang Guo, M. Maral Mouradian, Eileen White, Bing Xia

## Abstract

The PALB2 tumor suppressor plays key roles in DNA repair and has been implicated in redox homeostasis. Autophagy maintains mitochondrial quality, mitigates oxidative stress and suppresses neurodegeneration. Here we show that *Palb2* deletion in the mouse brain leads to motor deficits and that co-deletion of *Palb2* with the essential autophagy gene *Atg7* accelerates and exacerbates neurodegeneration induced by ATG7 loss. *Palb2* deletion leads to elevated DNA damage, oxidative stress and mitochondrial markers, especially in Purkinje cells, and co-deletion of *Palb2* and *Atg7* results in accelerated Purkinje cell loss. Further analyses suggest that the accelerated Purkinje cell loss and severe neurodegeneration in the double deletion mice are due to oxidative stress and mitochondrial dysfunction, rather than DNA damage, and partially dependent on p53 activity. Our studies uncover a role of PALB2 in mitochondrial regulation and a cooperation between PALB2 and ATG7/autophagy in maintaining redox and mitochondrial homeostasis essential for neuronal survival.

## Introduction

Neurodegenerative disorders affect approximately 15% of the population (Collaborators, 2019). Oxidative stress, which occurs when production of reactive oxygen species (ROS) in the cell exceeds the capacity of its antioxidant system, has been recognized as a major contributing factor in neurodegenerative diseases. ROS are primarily byproducts of aerobic metabolism during mitochondrial energy production (Forrester et al., 2018). ROS overproduction can lead to oxidative damage to both cellular and mitochondrial proteins, DNA, and membrane lipids, impairing a range of cellular functions, including the ability of mitochondria to synthesize ATP and carry out other metabolic functions (Murphy, 2009). The brain is particularly susceptible to oxidative stress due to its high levels of polyunsaturated fatty acids, which are prone to oxidation, and its relatively high metabolic rate and high oxygen demand (Singh et al., 2019; Watts et al., 2018), strict aerobic metabolism (Yang et al., 2014) and low levels of antioxidants (Baxter and Hardingham, 2016). ROS-induced damage to lipids, proteins and DNA is a common feature of all major neurodegenerative disorders (Butterfield et al., 2007; Canugovi et al., 2013).

One cellular mechanism that protects against neurodegeneration is autophagy. Autophagy is a cellular waste disposal and nutrient recycling mechanism that plays a key role in mitochondria quality control and metabolic rewiring upon stress (Ashrafi and Schwarz, 2013; Boya et al., 2018; Poillet-Perez and White, 2019), which can reduce ROS levels. Functional loss of autophagy causes accumulation of defective mitochondria along with other autophagy substrates and can lead to oxidative stress, cell death and senescence (Guo et al., 2016; Guo and White, 2013; Guo et al., 2013; Wang et al., 2016; Yang et al., 2020; Zong et al., 2016). Autophagy dysfunction is associated with neurological disorders such as Alzheimer’s disease, Parkinson’s disease, Huntington’s disease and amyotrophic lateral sclerosis (Li et al., 2008; Nixon et al., 2005; Ravikumar et al., 2004; Son et al., 2012; Vogiatzi et al., 2008). Neuron-specific ablation of essential autophagy genes *Atg5* or *Atg7* in mice results in motor and behavioral deficits, neurodegeneration, and greatly reduced survival (Hara et al., 2006; Komatsu et al., 2006).

While suppressing neurodegeneration, autophagy also has a role in promoting breast cancer (Wei et al., 2011). Approximately 5-10% of breast cancers are familial, resulting from inherited mutations in tumor suppressor genes such as *BRCA1*, *BRCA2* and *PALB2* (Yoshida, 2020). These genes encode large proteins that function in an BRCA1-PALB2-BRCA2 axis that is crucial to homologous recombination (HR)-mediated repair of DNA double strand breaks (DSBs) (Xia et al., 2006; Zhang et al., 2009). Moreover, we have shown that PALB2 promotes the accumulation and function of NRF2, a master antioxidant transcription factor, thereby promoting antioxidant response and reducing ROS levels (Ma et al., 2012). Interestingly, impaired autophagy, due to monoallelic loss of the essential autophagy gene *Becn1*, delayed and reduced *Palb2*-associated mammary tumorigenesis, suggesting that autophagy facilitates mammary tumor development following the loss of PALB2 (Huo et al., 2013).

In the current study, we aimed to co-delete *Palb2* and *Atg7* in the mammary gland using Cre recombinase driven by the promoter of whey acidic protein (*Wap-Cre*), to further study the role of autophagy in PALB2-assciated breast cancer. Unexpectedly, *Wap-Cre* caused efficient deletion of the genes in the brain, and co-deletion of the genes led to accelerated and more severe neurodegeneration compared with *Atg7* deletion alone. This was further supported by findings from mice with whole-body deletion of *Palb2* and *Atg7* driven by *Ubc-Cre-ERT2*. Detailed analyses of the brain revealed a novel role of PALB2 in regulating mitochondrial homeostasis and that the double deletion brain sustained more oxidative stress, rather than DNA damage, than either single deletion brain. Moreover, p53 was found to be induced by both *Palb2* and *Atg7* deletions, and further deletion of *Trp53* significantly but partially rescued the neurodegenerative phenotype of *Palb2*/*Atg7* double deletion mice. Our studies uncover a novel function of PALB2 and underscore the important roles and complex interplay of oxidative stress, autophagy, mitochondria and p53 in brain neurodegeneration.

## Results

### Survival, tumor development and motor behavioral deficits of mice with individual and combined ablation of *Palb2* and *Atg7*

As heterozygous disruption of *Becn1* delays *Palb2*-associated mammary tumor development (Huo et al., 2013), we sought to further examine the role of autophagy in mammary tumor development following PALB2 loss. *Palb2^f/f^*;*Wap-Cre* (*Palb2*-CKO) mice were crossed with *Atg7*-floxed mice to generate mice with single and combined knockout of the two genes in the mammary gland. *Palb2*-CKO mice showed a median overall survival (T_50_) of 649 days (Figure 1A), and most of these mice developed spontaneous tumors (Figure 1B). Notably, only 6/19 (32%) of *Palb2*-CKO mice developed mammary tumors (Figure 1C), likely due to a mostly C57BL/6 genetic background, which is known to be more resistant to mammary tumorigenesis. *Atg7^f/f^*;*Wap-Cre* (*Atg7*-CKO) mice had slightly shorter overall survival (T_50_=604 days) (Figure 1A); 1 of the 22 mice (4.5%) developed mammary tumor and 9 (45.5%) developed other tumors (Figure 1C).

**Figure 1.**
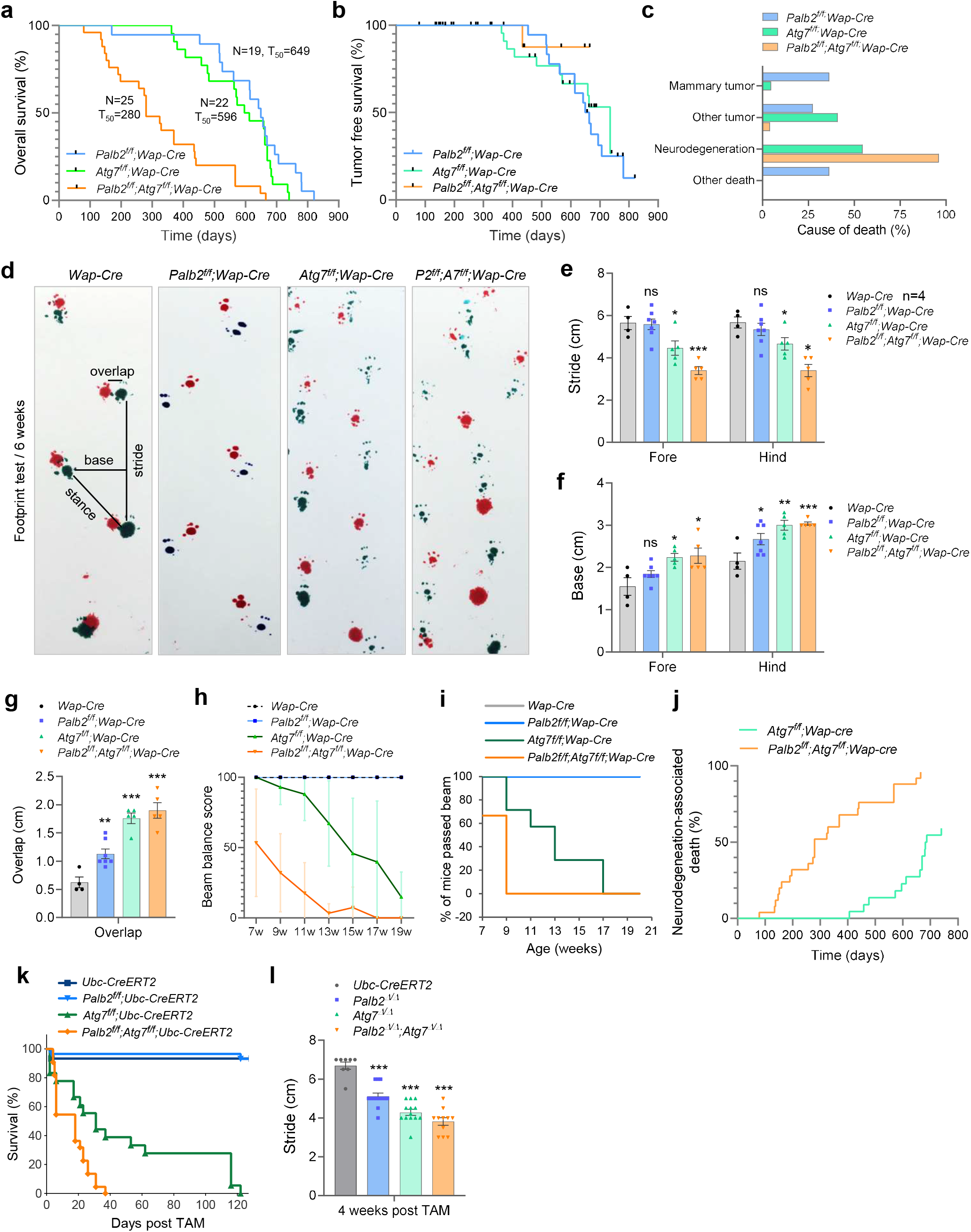
Survival curves and motor deficits of *Palb2* and *Atg7* deletion mice. (A-C) Overall survival (A), tumor-free survival (B) and observed causes of death (C) of *Wap-Cre* driven CKO mice. Black dots in b indicate censoring events (mice that died without tumor). (D-G) Footprint test of the CKO mice. Representative footprints and variables measured are shown in D, and results of stride, base and overlap measurements are shown in E, F and G, respectively. n=4-7. (H and I) Beam balance test of the CKO mice. Beam balance scores are shown in H and the percentage of mice that passed the beam in I. n=4-7. (J) Neurodegeneration-associated death in the CKO cohorts. (K-I) Survival curves (K) and stride length (I) of mice with *Ubc-Cre-ERT2* driven, whole body knockout (WBKO) of *Palb2* and/or *Atg7*. n=8-13. Error bars indicate standard errors (SE). ns, not significant; * p≤0.05; ** p≤0.01; *** p≤0.001.

Unexpectedly, *Atg7*-CKO mice showed progressive motor deficits, manifesting primarily as gait ataxia suggestive of cerebellar involvement. Note that although the *Wap-cre* used here was designed to expresses Cre specifically in the mammary gland, Cre activity has been reported in the brain (Wagner et al., 1997). Gait analysis at 6 weeks of age revealed an ataxic walking pattern of *Atg7*-CKO mice (Figure 1D). Compared with *Wap-Cre* (control) mice, mean stride length of *Atg7*-CKO mice was significantly decreased (Figure 1E), and both mean base width and overlap were significantly increased (Figures 1F and 1G). Beam balance test also showed gradual impairment of motor coordination in *Atg7*-CKO mice (Figures 1H, and 1I). Additionally, these mice showed limb clasping reflexes when suspended by the tails (data not shown). These motor behavioral phenotypes are consistent with previous reports that loss of *Atg7* in the brain leads to neurodegeneration (Inoue et al., 2012; Komatsu et al., 2007; Yang et al., 2020). The mice were able to survive with the motor deficits until approximately 400 days of age and then started to die. Eventually, 12/22 (54.5%) of the mice succumbed to neurodegeneration-associated death (Figure 1J), defined as death with severe motor deficits and without tumor or other overt pathological conditions seen upon dissection.

Remarkably, *Palb2^f/f^*;*Atg7^f/f^*;*Wap-Cre* (*Palb2*;*Atg7*-CKO) mice exhibited greatly reduced overall survival (T_50_=280 days) (Figure 1A), even though their tumor development was also greatly reduced (Figure 1B). These double CKO mice showed more severe motor deficits, specially in terms of stride length and balance (Figures 1E, 1H and 1I), and died of motor deficits much earlier than *Atg7*-CKO mice (Figure 1J). Only 1 out of 25 (4%) of these mice developed cancer, and all the rest died from severe motor deficits. The more severe motor deficits of the double CKO mice and the striking acceleration of death associated with neurodegeneration in these mice suggest that the two genes may synergistically suppress neurodegeneration. Although *Palb2-*CKO mice did not show any overt behavioral deficits, footprint test showed increased hind base width and overlap (Figures 1F and 1G), suggestive of mild motor dysfunction.

We also crossed *Atg7*- and *Palb2*-floxed mice with mice carrying a *Ubc-Cre-ERT2* allele, which allow for tamoxifen (TAM)-induced whole-body knockout (WBKO) of floxed genes (Ruzankina et al., 2007). One week after (5 daily) TAM injections, efficient deletions of both *Palb2* and *Atg7* were observed in brain tissues (Figure S1A). The resulting *Palb2*-WBKO (*Palb2*^Δ/Δ^) mice appeared normal and survived for at least a year. In contrast, a small number of *Atg7*^Δ/Δ^ mice died from toxicity and/or infections within the first week, the rest soon manifested motor deficits that worsened progressively over time, and the mice died within 120 days (Figure 1K), as we have shown recently (Karsli-Uzunbas et al., 2014; Yang et al., 2020). Notably, *Palb2*^Δ/Δ^;*Atg7*^Δ/Δ^ mice had even shorter survival compared with *Atg7*^Δ/Δ^ mice (T_50_=16 vs 30 days, p<0.05). Gait analysis at 4 weeks post TAM treatment showed that both *Palb2*^Δ/Δ^ mice and *Atg7*^Δ/Δ^ mice had significantly shortened stride length, and the length was even shorter in *Palb2*^Δ/Δ^;*Atg7*^Δ/Δ^ mice (Figure 1L). These double knockout mice also died primarily from severe motor deficits, except those that died shortly after injection due to toxicity and/or infections. Thus, loss of PALB2 in the brain leads to mild motor impairment but greatly exacerbates neurological phenotype induced by *Atg7* deficiency.

### Loss of PALB2 exacerbates neurodegeneration induced by ATG7 loss

To better understand the cause of the neurological phenotype in our models, we assessed the “leaky” expression of Cre in multiple tissues. Indeed, deletion of *Palb2* was observed in the brains of *Palb2*-CKO and *Palb2*;*Atg7*-CKO mice (Figures S1B and S1C). Additionally, we analyzed the protein level of p62/SQSTM1, an autophagy adaptor protein that accumulates upon autophagy inhibition (Bjorkoy et al., 2005), in the whole brain by immunofluorescence (IF). In *Atg7*-CKO mice, aberrant accumulation of p62 was detected throughout the brain, with intense staining signals in brain stem and scattered signals in cerebral cortex and cerebellum (Figure 2A, green). Western blotting further confirmed strong p62 accumulation in midbrain, cerebellum and cerebral cortex (Figure S1D). We also stained for tyrosine hydrolase (TH), a marker of dopaminergic neurons in the substantia nigra that degenerate in Parkinson’s disease. However, no difference in TH staining was observed between control and any of the CKO mice (Fig. 2a, red), indicating that the dopaminergic neurons were likely intact by 10 weeks of age, consistent with a previous report (Friedman et al., 2012).

**Figure 2.**
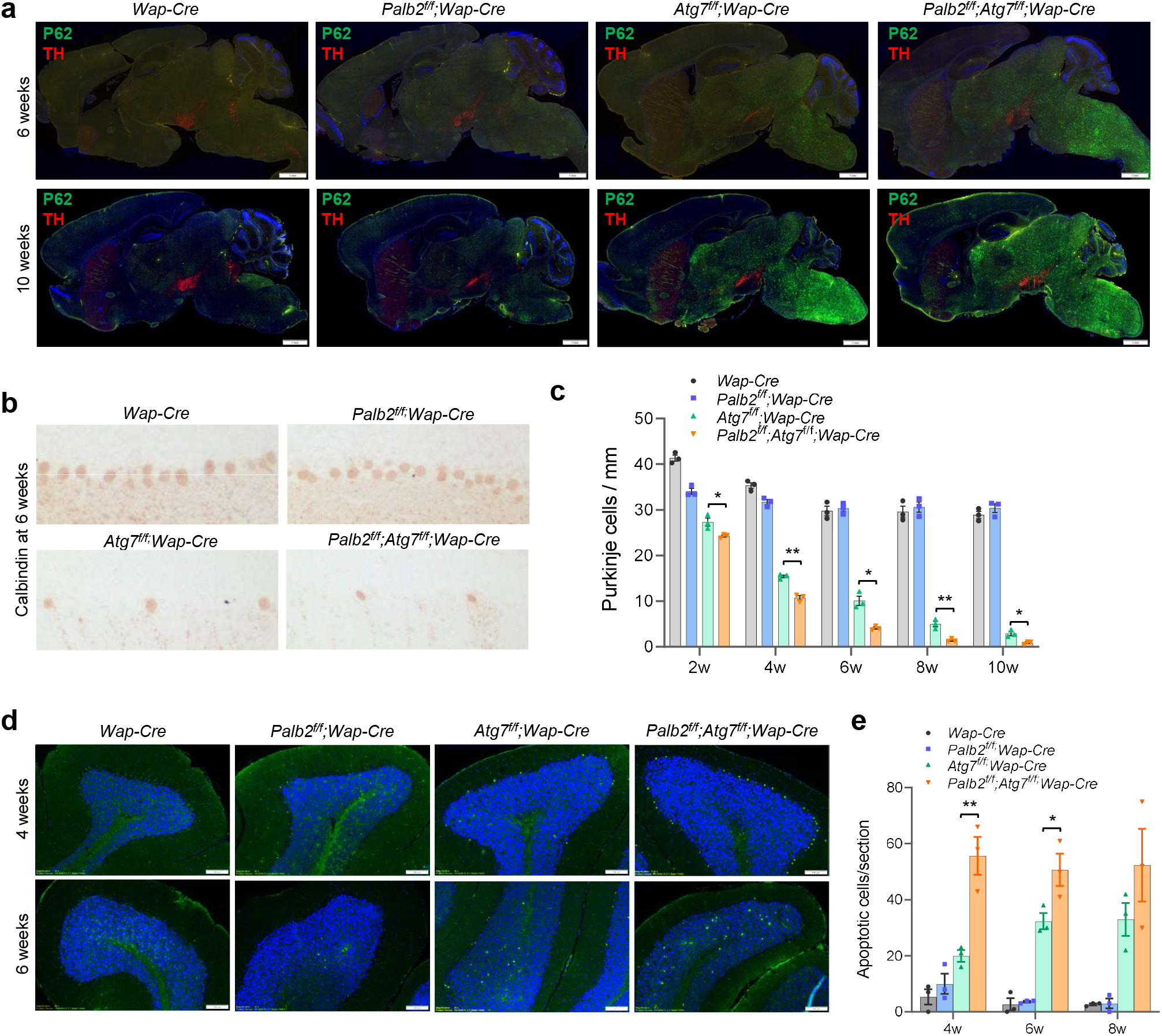
Neuronal degeneration in *Palb2* and *Atg7* CKO mice. (A) Representative immunofluorscence staining of p62/SQSTM1 (green) and TH (red) in the midsagittal section of the brain. Nuclei were stained with DAPI. (B and C) Representative IHC staining of cerebellum with Purkinje cell marker Calbindin (B) and quantification of Purkinje cells (C) at the indicated weeks of age. n=3. (D and E) Representative TUNEL images (D) and quantification of positive cells (E) in the brains of CKO mice at different weeks of age. n=3. Error bars indicate SE. * p≤0.05; ** p≤0.01.

It is well know that *Atg7* deletion in the brain leads to loss of Purkinje cells in the cerebellum (Komatsu et al., 2006; Komatsu et al., 2007), which are primary controller of motor coordination and balance, and the motor deficits observed in the present study are consistent with degeneration of these neurons. Indeed, immunohistochemistry (IHC) for Calbindin, a marker for Purkinje cells, demonstrated progressive loss of these cells in *Atg7*-CKO mice between 2 and 10 weeks of age (Figures 2B and 2C). *Palb2*-CKO mice showed only a modest reduction in the number of these cells at 2 weeks but no further reduction later, while the double CKO mice showed more pronounced and significantly accelerated Purkinje cell loss compared with *Atg7*-CKO mice (Figure 2C). Apoptotic cells were detected in both Purkinje cell layer and granular cell layer of both *Atg7*-CKO and *Palb2;Atg7*-CKO mice, with the double CKO mice showing more apoptotic Purkinje cells than *Atg7*-CKO mice at 4 weeks of age (Figure 2D). At 6 weeks, while *Atg7*-CKO mice still had some apoptotic Purkinje cells, these cells were hardly detectable in *Palb2;Atg7*-CKO mice. At this age, the double CKO mice also had significantly more apoptotic cells in the granular cell layer than did *Atg7*-CKO mice (Figures 2D and 2E). Collectively, the above observations indicate that loss of *Palb2* exacerbates and accelerates the neurodegeneration induced by *Atg7* deletion.

### DNA damage and oxidative stress in *Palb2* and *Atg7* deleted brains

To begin to understand the cause of Purkinje cell loss and the neurodegenerative phenotype of our mice, and considering the known roles of PALB2 and autophagy in DNA repair and redox homeostasis, we assessed the levels of γH2AX, a marker for DSBs, 8-oxo-deoxyguanidine (8-oxo-dG), a marker for oxidative DNA damage, and 4-HNE, a product of lipid peroxidation, in the cerebellum of *Palb2* and *Atg7* CKO mice by IHC.

In both control and the CKO mice, positive γH2AX signals were only observed in Purkinje cells (Figure 3A). Unlike the typical DNA damage-induced γH2AX nuclear foci, γH2AX in Purkinje cells was observed mostly in a single large and round structure in the nucleus, whose nature remains to be determined. In control mice, 15-30% of Purkinje cells showed positive γH2AX staining, whereas this percentage was always higher than 50% in *Palb2*-CKO mice (Figure 3B). In *Atg7*-CKO mice, the percentage of γH2AX positive Purkinje cells was comparable to that in *Wap-Cre* control mice at 2 and 4 weeks of age; at 6 weeks and beyond, most of the Purkinje cells had been lost, and the remaining cells showed higher γH2AX positivity. In the double CKO mice, γH2AX positivity in Purkinje cells was similar to that in *Palb2* single CKO mice at 2 and 4 weeks of age; at 6 and 10 weeks of age, the remaining few Purkinje cells in the double CKO mice showed even higher positivity than those in *Palb2* single CKO mice. Note that the high γH2AX positivity in the *Atg7*-CKO and *Palb2*;*Atg7*-CKO Purkinje cells at later time points cannot be interpreted with confidence due to cell shrinkage and possible ongoing apoptosis.

**Figure 3.**
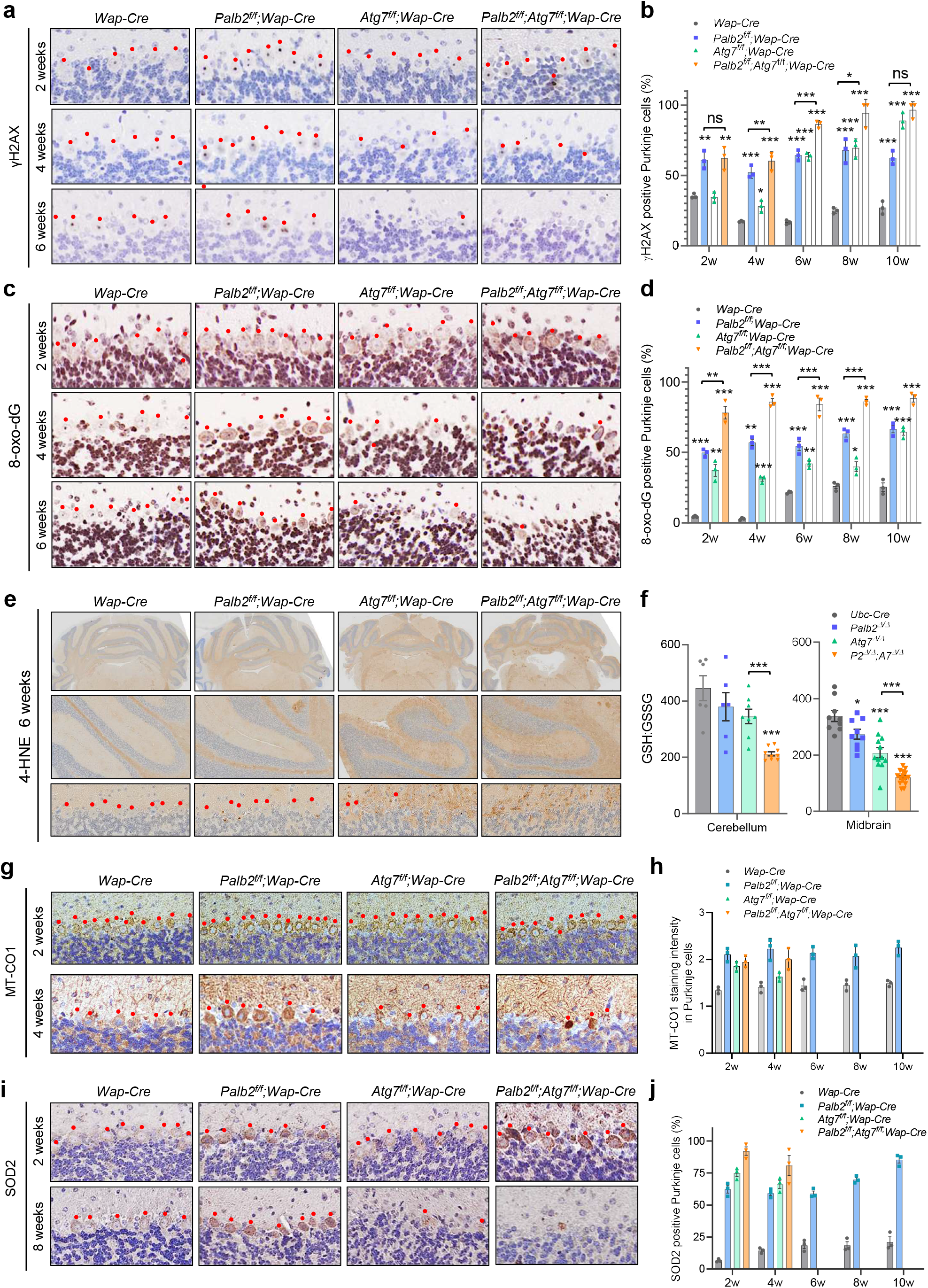
Increased DNA damage and oxidative stress and mitochondrial markers in Purkinje cells of *Palb2* and *Atg7* deletion mice. (A-D) Representative IHC images and quantification of DNA DSB marker γH2AX (A and B) and oxidative DNA damage marker 8-oxo-dG (C and D) in Purkinje cells of *Wap-Cre* driven CKO mice at 2, 4, 6, 8 and 10 weeks. n=3. Red dots indicate Purkinje cells. (E) Representative IHC staining of lipid peroxidation by-product 4-HNE in the CKO mice at 6 weeks of age. (F) GSH:GSSG ratios in the cerebellum and midbrain of *Ubc-Cre-ERT2* driven WBKO mice. GSH and GSSG were measured by LC–MS/MS at 4 weeks after tamoxifen injection. n=6-9 for cerebellum and n=9-15 for midbrain. (G-J) Representative IHC images and quantification of MT-CO1 (G and H) and SOD2 (I an J) in Purkinje cells of CKO mice at 2, 4, 6, 8 and 10 weeks. Red dots indicate Purkinje cells. n=3. Error bars indicate SE. * p≤0.05; ** p≤0.01; *** p≤0.001.

γH2AX was also analyzed in the brains of the afore-described WBKO (Δ/Δ) mice. As expected, increased γH2AX signal was found in Purkinje cells of *Palb2*^Δ/Δ^ mice at 4 weeks after tamoxifen injection (Figure S2A). At this time, Purkinje cells had been mostly lost in *Atg7*^Δ/Δ^and *Palb2*^Δ/Δ^*;Atg7*^Δ/Δ^ mice; however, increased γH2AX was detected in a subset of granular cells in these mice, suggesting that ATG7 plays a role, directly or indirectly, in DNA damage repair, consistent with our recent findings (Yang et al., 2020). Moreover, we subjected the mice to 10 Gy of whole-body gama radiation. As expected, *Palb2*^Δ/Δ^ mice showed substantially reduced survival compared with either control or *Atg7*^Δ/Δ^ mice (Figure S2B). Notably, however, *Palb2*^Δ/Δ^*;Atg7*^Δ/Δ^ mice exhibited similar survival to that of *Palb2*^Δ/Δ^ mice post radiation, suggesting that the DNA repair deficit of *Palb2*^Δ/Δ^*;Atg7*^Δ/Δ^ mice stem primarily from PALB2 deficiency.

8-oxo-dG staining signals were observed in both Purkinje and granular cells, but any clear difference between control and CKO mice was mainly found in Purkinje cells (Figures 3C, d). At 2 weeks of age, very few Purkinje cells in the control mice showed clear positive 8-oxo-dG staining and the signals were also weak, whereas more cells in *Palb2*-CKO and *Atg7*-CKO mice were positive and the signals were slightly stronger. Notably, *Palb2*;*Atg7-*CKO mice had even more 8-oxo-dG positive Purkinje cells at this time point, and the signals were also substantially stronger. At 4 weeks, the staining in *Palb2*-CKO mice was much stronger than that at 2 weeks and also much stronger than that in either control or *Atg7*-CKO mice, although *Atg7*-CKO mice showed slightly higher staining intensity than the control. At 6 weeks and beyond, Purkinje cell 8-oxo-dG staining in *Palb2*-CKO mice remained much stronger than that in control mice. These results suggest that PALB2 is critical for preventing DNA oxidation in Purkinje cells, whereas ATG7 plays a less prominent role. On the other hand, 4-HNE staining was evidently more intense in *Atg7*-CKO and *Palb2*;*Atg7*-CKO brains (Figure 3E). Finally, we also determined the ratio of reduced to oxidized glutathione (GSH:GSSG), a key indicator of cellular redox state (Aquilano et al., 2014), in the brains of the WBKO mice. The ratio was decreased in *Palb2*^Δ/Δ^ and *Atg7*^Δ/Δ^ mice and further decreased in *Palb2*^Δ/Δ^*;Atg7*^Δ/Δ^ mice (Figure 3F). Taken together, these results reveal both similar and distinct impacts of PALB2 and ATG7 loss on DNA damage and oxidative stress in the brain.

### Requirement of PALB2 for mitochondrial homeostasis in the brain

Given the key role of autophagy in mitochondrial quality control and mitochondrial dysfunction in neurodegenerative diseases, we analyzed two mitochondria markers by IHC. First, we examined MT-CO1, a subunit of respiratory complex IV (Rak et al., 2016). At 2 weeks of age, Purkinje cells in both *Palb2*-CKO and *Atg7*-CKO mice appeared to have slightly higher levels of MT-CO1 than control; at 4 weeks, MT-CO1 signals in *Palb2*-CKO Purkinje cells were much stronger than control, while signals in *Atg7*-CKO and *Palb2;Atg7* double CKO cells were condensed presumably due to cell shrinkage (Figures 3G and 3H). Next, we stained for SOD2 (MnSOD), a superoxide dismutase that is localized in the mitochondrial intermembrane space and converts superoxide anion resulting from leaked electrons from the electron transport chain to hydrogen peroxide (H_2_O_2_) (Kokoszka et al., 2001). At 2 weeks, *Palb2*-CKO Purkinje cells showed substantially stronger SOD2 staining than controls, signals in *Atg7*-CKO cells were similar to, if not slightly stronger than, controls, while the cells in the double CKO mice showed strongest SOD2 staining (Figures 3I and 3J). At 8 weeks, SOD2 expression remained much higher in *Palb2*-CKO Purkinje cells. Additionally, increased MT-CO1 was also found in Purkinje cells of *Palb2*^Δ/Δ^ mice (Figure S2A).

### Mitochondrial abnormality, oxidative stress, and cell death upon combined loss of PALB2 and ATG7 in human medulloblastoma cells

To confirm the above findings in human cells, we used CRISPR/Cas9 to knock out *PALB2* and *ATG7* in the DAOY medulloblastoma cells, which have a cerebellar origin (Jacobsen et al., 1985). *PALB2*;*ATG7* double knockout (DKO) clones were then obtained by further knocking out *ATG7* in a sequence validated and functionally characterized *PALB2*-KO clone. Consistent with the essential role of ATG7 in LC3 lipidation and autophagosome formation(Klionsky and Schulman, 2014), *ATG7*-KO cells showed reduced LC3-I o LC3-II conversion and increased p62 accumulation (Figure 4A). Consistent with our in vivo findings, MT-CO1 and MT-CO2 were increased in *PALB2*-KO, *ATG7*-KO and *PALB2*;*ATG7*-DKO cells, implying increased mitochondrial mass upon loss of either PALB2 or ATG7. Increase in SOD2 abundance was less consistent but was seen in *PALB2*-KO cells most of the time and in *ATG7*-KO cells sometimes, suggesting that it might be dependent on cell growth condition or state. *ATG7*-KO cells grew at a rate similar to that of control cells, while *PALB2*-KO cells showed significantly decreased growth rate, and a further decrease was seen for *PALB2;ATG7-*DKO cells (Figure 4B). Both *PALB2*-KO and *ATG7*-KO cells showed moderate boundary shrinkage, and the DKO cells showed overt boundary shrinkage along with soma cavitation (Figure 4C). Spontaneous cell death, including both apoptosis and necrosis, was increased in *PALB2-*KO and *ATG7*-KO cells and further increased in the DKO cells (Figures 4D and 4E).

**Figure 4.**
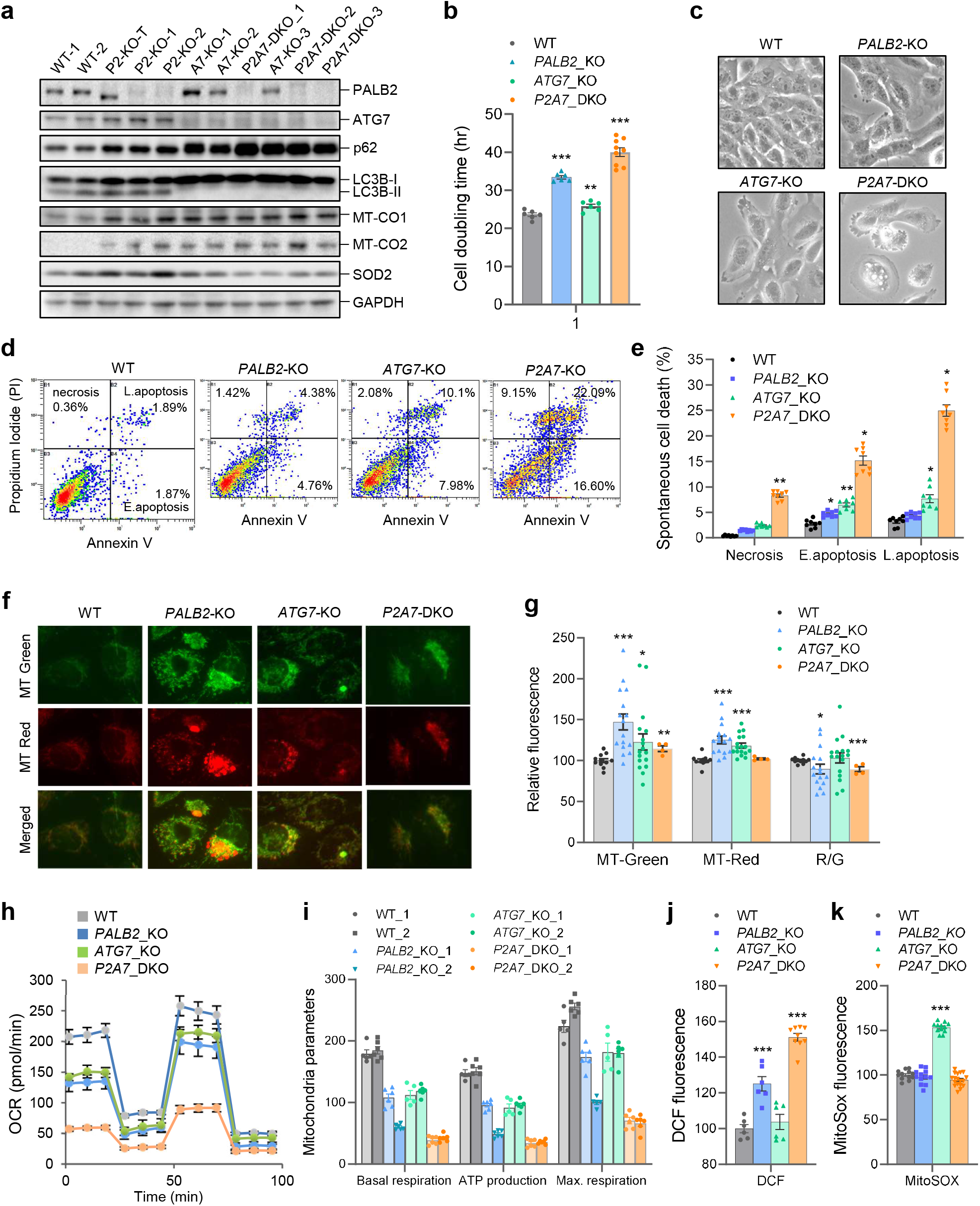
Mitochondrial abnormality, oxidative stress, and apoptosis in *PALB2* and *ATG7* KO human medulloblastoma cells. (A) Western blot analysis of whole cell homogenates of WT and KO cells. All cell lines were from single clones. The WT lines were false positive clones from the *PALB2* CRISPR/Cas9 KO procedure. (B) Doubling times of WT and KO cells. (C) Representative bright field microscopy images of the cells. (D and E) Representative flow cytometry plots (d) and quantification (e) of Annexin V assay analysis of the cells. (F and G) Mitochondrial analyses of WT and KO cells with MitoTracker-Green and MitoTracker-Red. Representative images of stained cells are shown in f and FACS-based detection/quantification of the signals in g. Avg. R/G, average ratios of Red vs Green signals (in percentage). (H) Oxygen consumption rate (OCR) in WT and KO cells as measured by the Seahorse assay. (I) Bar graph of mitochondrial functional parameters as measured in panel h. (J and K) Levels of reactive oxygen species in WT and KO cells as measured by DCF (J) and MitoSOX (K) staining followed by FACS-based detection and quantification. Error bars indicate SE. * p≤0.05; ** p≤0.01; *** p≤0.001.

Next, we analyzed mitochondria using MitoTracker Green and MitoTracker Red, which measure mitochondrial mass and membrane potential, respectively (Figure 4F). *ATG7*-KO cells showed an increase in total mitochondrial mass, consistent with previous reports (Karsli-Uzunbas et al., 2014; Kimmelman and White, 2017; Strohecker et al., 2013). Notably, an even larger increase in mitochondrial mass was observed in *PALB2*-KO cells. Both single KO cells showed modestly increased membrane potential compared with WT cells, whereas the DKO cells showed no change. The ratio of MitoTracker Red to Green signals (R/G) of *PALB2*-KO, *ATG7*-KO and the DKO cells was 85.09%, 92.82% and 88.76%, respectively, of wt level, indicative of impaired mitochondrial quality in all 3 cell types. The increased mitochondrial mass in *PALB2*-KO cells corroborates with our in vivo findings (Figure 3) and lends further support to the notion that loss of PALB2 leads to increased mitochondrial biogenesis, while the decreased R/G ratio in the cells suggests lower overall mitochondria quality or a defect in the clearance of damaged mitochondria. To evaluate if mitochondrial respiration was affected by *PALB2* and/or *ATG7* inactivation, oxygen consumption rate (OCR) was measured by the Seahorse assay. Basal mitochondrial respiration, ATP-linked respiration and maximal repiration were moderately decreased in both *PALB2*-KO and *ATG7*-KO cells, but dramatically decreased in *PALB2;ATG7*-DKO cells (Figures 4H and 4I). These data suggest that a strong defect in mitochondrial respiration may be a major contributor to the spontaneous death of DKO cells.

Mitochondria are both a major producer and a primary target of ROS (Zorov et al., 2014). Consistent with our in vivo observations, cellular ROS level measured by DCF, which mainly detects H_2_O_2_, was significantly higher in *PALB2*-KO cells and even higher in *PALB2;ATG7*-DKO cells; however, no change in DCF readout was observed in *ATG7*-KO cells (Figure 4J). In contrast, mitochondrial superoxide, as measured by MitoSOX, was increased in *ATG7*-KO cells but not in *PALB2*-KO or *PALB2;ATG7*-DKO cells (Figure 4K). The increased superoxide level in *ATG7*-KO cells is consistent with recent reports (Nahapetyan et al., 2019; Trentesaux et al., 2020), and the lack of any change in DCF readings suggests that the overproduced superoxide in these cells is not converted to H_2_O_2_ in a timely fashion, which could lead to mitochondrial membrane oxidation. On the other hand, the increase in DCF signals in *PALB2*-KO cells without any concomitant increase in superoxide suggests that the higher ROS level may not originate from mitochondria; another possibility is that mitochondria in *PALB2*-KO cells indeed overproduce superoxide, but the extra superoxide is quickly dismutated into H_2_O_2_ by the increased SOD2, leading to higher DCF signals. These data are largely consistent with our findings from the mouse brain and support the notion that a combination of abnormal mitochondria function and increased oxidative stress leads to apoptosis in Purkinje cells and certain other neurons, hence the motor deficits.

### Loss of BRCA2 delays the onset of neurodegeneration induced by ATG7 deficiency

To assess whether an HR deficiency and increased DSB formation were a significant cause of the more severe neurodegeneration in *Palb2*;*Atg7*-CKO mice than in *Atg7*-CKO mice, we generated *Brca2^f/f^;Atg7^f/f^;Wap-cre* (*Brca2*;*Atg7*-CKO) mice. Interestingly, unlike *Palb2*;*Atg7*-CKO mice, *Brca2*;*Atg7*-CKO mice had similar overall survival and neurodegeneration-associated death as those of *Atg7*-CKO mice (Figures 5A and 5B). Also unlike *Palb2*, co-deletion of *Brca2* with *Atg7* did not exacerbate the neurological phenotype of *Atg7*-CKO mice; instead, it significantly moderated Purkinje cell loss caused by *Atg7* deficiency from 2 to 10 weeks of age (Figure 5C). Both gait analysis and beam balance test also revealed a rescue of motor coordination and balance defects by co-deletion of *Brca2* with *Atg7* (Figures 5D and 5E).

**Figure 5.**
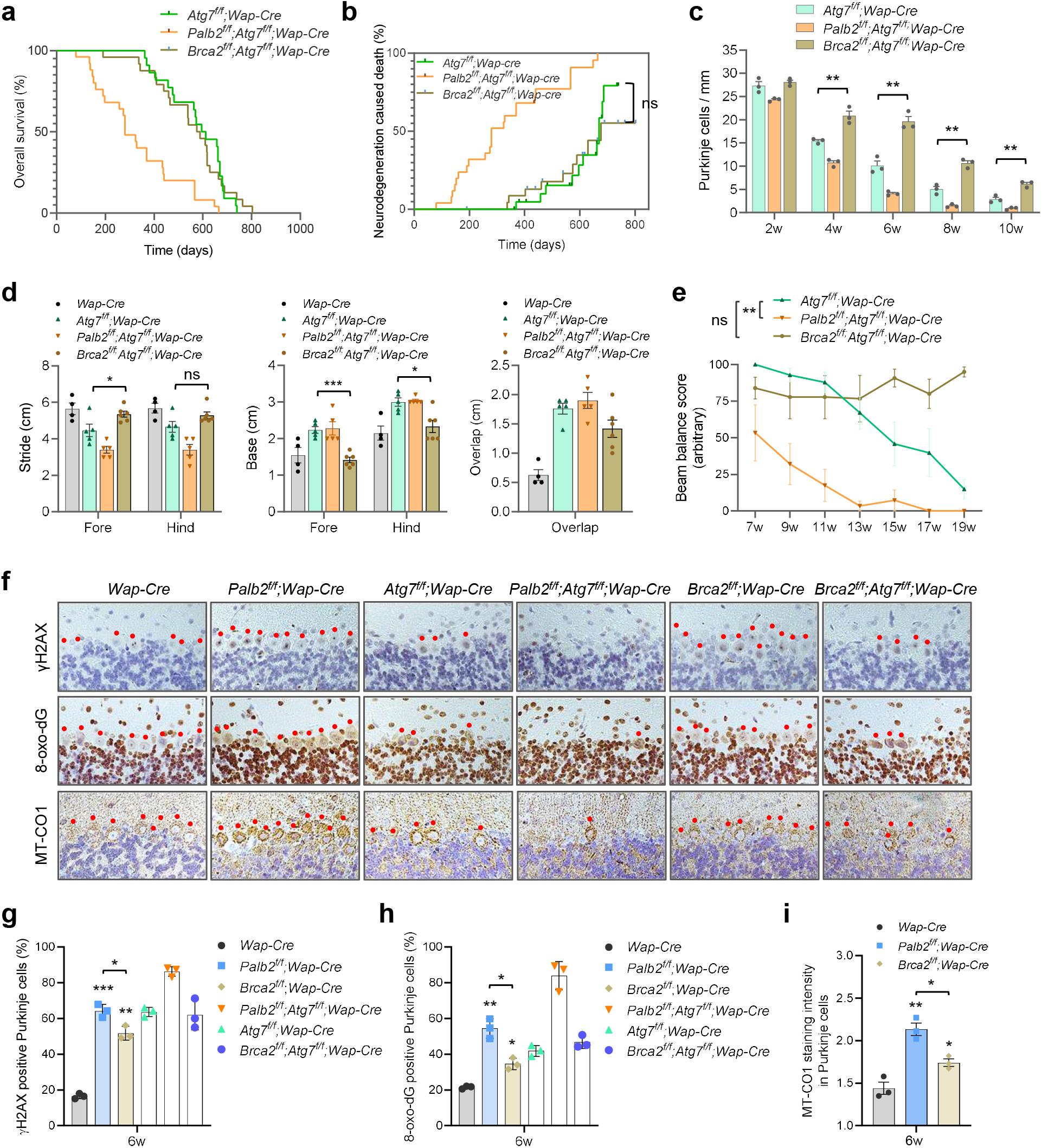
Comparative analyses of the impacts of *Brca2* and *Palb2* deletions in wt and *Atg7*-CKO mice. (A and B) Overall survival (A) and neurodegeneration-associateded death (B) of CKO mice. (C) Purkinje cell numbers in CKO mice at 2 to 10 weeks of age (n=3). (D) Footprint measurements of 6 weeks old CKO mice (n=5-7). (E) Beam balance scores of CKO mice at different ages (n=5-7 mice per time point). (F) Representative images of Calbindin, γH2AX, 8-oxo-dG and MT-CO1 IHC in the cerebellum of 6 weeks old CKO mice. (G-I) Quantification of γH2AX (G), 8-oxo-dG (H) and MT-CO1 (I) staining positivity in Purkinje cells (n=3). Error bars indicate SE. * p≤0.05; ** p≤0.01; *** p≤0.001.

To understand why ablation of *Brca2* produced a different effect from that of *Palb2* ablation, we compared γH2AX, 8-oxo-dG and MT-CO1 in the brains of the mice at 6 weeks of age. As shown in Figures 5F-I, γH2AX positivity in Purkinje cells of *Brca2*-CKO mice was slightly lower than that in *Palb2*-CKO mice but still much higher than that in control mice; in contrast, their 8-oxo-dG and MT-CO1 staining signals were substantially weaker than those in *Palb2*-deleted Purkinje cells. The modestly higher γH2AX positivity in *Palb2*-deleted Purkinje cells compared with *Brca2*-deleted cells may be attributed to higher oxidative stress, which can cause DSBs. Compared with the remaining Purkinje cells in *Atg7*-CKO mice, the same cells in *Brca2*;*Atg7*-CKO mice showed similar γH2AX positivity. The staining patterns of both MT-CO1 and 8-oxo-dG in Purkinje cells of *Brca2*;*Atg7*-CKO mice were also similar to that in *Atg7-*CKO mice. Therefore, loss of DNA repair function of PALB2 is unlikely to be a significant cause of the more severe neurological phenotype of *Palb2*;*Atg7*-CKO mice, and that increased oxidative stress and/or mitochondrial dysfunction may underlie this exacerbated phenotype.

### Loss of p53 delays Purkinje cell death and prolongs the survival of *Palb2;Atg7*-CKO mice

Our recent study showed that ATG7 limits p53 activation and p53-induced neurodegeneration (Yang et al., 2020). To determine whether p53 played a role in the shortened survival of *Palb2;Atg7*-CKO mice, we set up cohorts of these mice either without or with floxed *Trp53*. Compared with *Palb2;Atg7*-CKO mice, overall survival of *Palb2;Atg7;Trp53-*CKO mice was significantly prolonged (T_50_=448 vs 280 days, p=0.0165) (Figure 6A), and neurodegeneration-associated death was greatly reduced and delayed (Figure 6B). Moreover, inactivation of *Trp53* combined with prolonged survival allowed for efficient tumor development (Figure 6C). While 95% of *Palb2;Atg7*-CKO mice died from neurodegeneration, 45% of *Palb2;Atg7;Trp53*-CKO mice developed mammary tumors and 21% developed other tumors, with neurodegeneration causing the death of only 30% of the mice.

**Figure 6.**
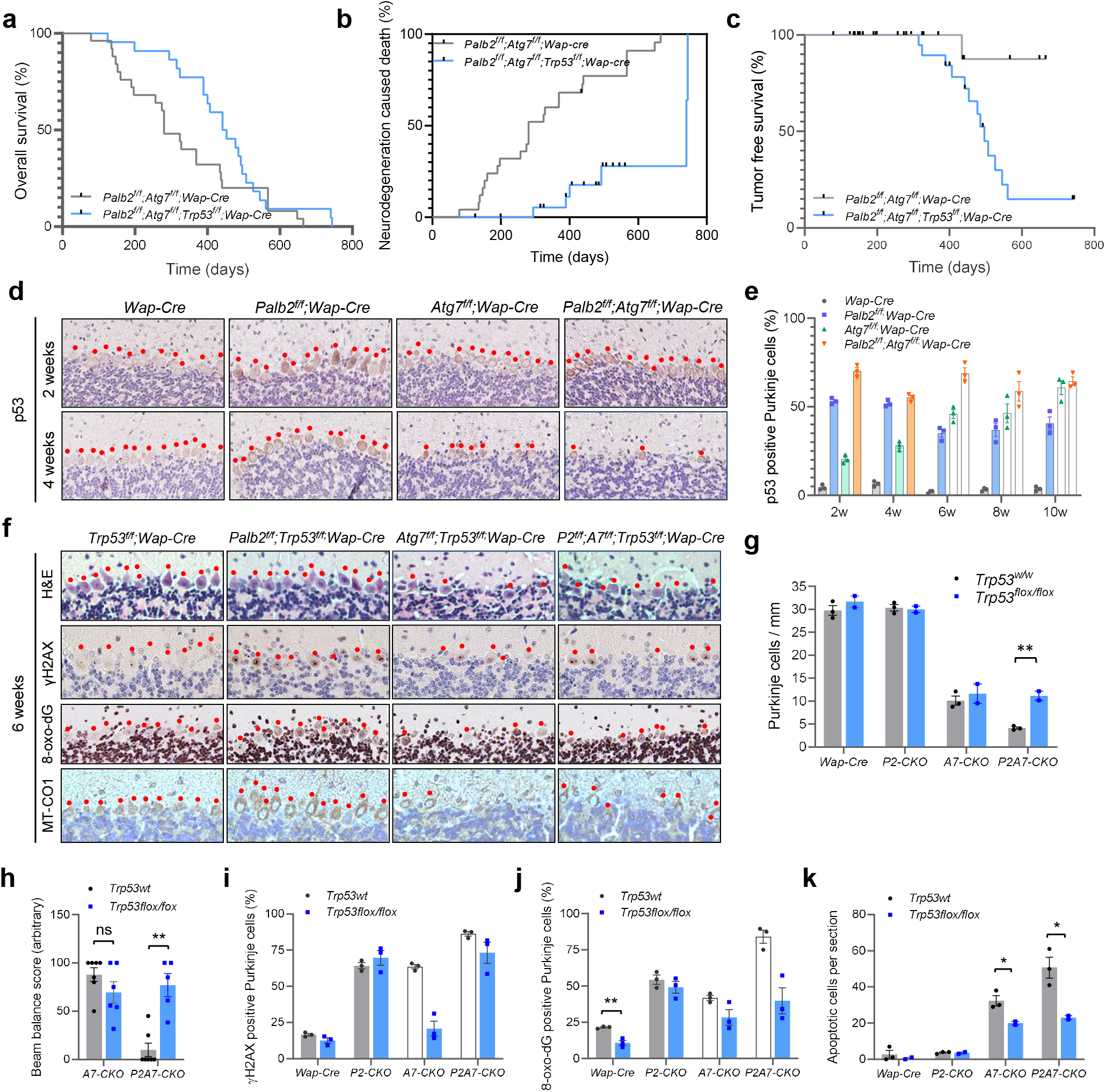
Role of p53 in Purkinje cell death and the survival of *Palb2;Atg7*-CKO mice. (A-C) Overall survival (A), neurodegeneration-associated death (B) and tumor free survival (C) of *Palb2*-CKO and *Palb2;Atg7*-CKO mice without and with *Trp53* co-deletion. Where applicable, black dots indicate the censoring events. (D and E) Representative p53 IHC images (D) and quantification of p53 positive Purkinje cells (E) in the cerebellum of control (*Wap-Cre*) and CKO mice at different ages (n=3 per time point). (F) Representative H&E and IHC images of γH2AX, 8-oxo-dG and MT-CO1 in the cerebellum of CKO mice at 6 weeks of age. (G) Density of Purkinje cells in CKO mice at 6 weeks of age (n=2-3). (H) Beam balance scores of CKO mice at 11 weeks of age (n=5-7). (I and J) Quantification of γH2AX positive (I) and 8-oxo-dG positive (J) Purkinje cells at 6 weeks of age (n=2-3). (K) Number of apoptotic cells in the cerebellum of 6 weeks old control and CKO mice (n=2-3). Error bars indicate SE. ns, not significant; * p≤0.05; ** p≤0.01.

The above results suggest that p53 was activated upon loss of PALB2 and/or ATG7 in the brain which subsequently led to neuronal apoptosis. Indeed, IHC analysis revealed that at 2 weeks of age, p53 protein level was moderately increased in *Atg7*-CKO Purkinje cells in terms of both staining intensity and number of positive cells, and even higher p53 accumulation was seen in Purkinje cells of *Palb2*-CKO and the double CKO mice (Figures 6D and 6E). The overall situation was similar at 4 and 6 weeks, except that some or most Purkinje cells in *Atg7*-KO mice and *Palb2*;*Atg7*-CKO mice had been lost.

Co-deletion of *Trp53* produced no significant effect on Purkinje cell numbers in control, *Palb2*-CKO or *Atg7*-CKO mice; however, the severe loss of Purkinje cells in *Palb2*;*Atg7*-CKO mice was significantly rescued (Figures 6F and 6G). Interstingly, when *Trp53* was co-deleted, Purkinje cell number in *Palb2;Atg7*-CKO mice was restored to the same level as that in *Atg7*-CKO mice (Figure 6G), suggesting that the severe neurodegenerative phenotype in *Palb2*;*Atg7*-CKO mice is likely an *Atg7* null phenotype exacerbated by a further induction of p53 elicited by loss of PALB2. Consistent with the rescue of Purkinje cell number, motor function of *Palb2;Atg7;Trp53*-CKO mice was substantially preserved, with the mice achieving similar beam balance scores as *Atg7*-CKO mice at 11 weeks of age (Figure 6H).

Deletion of *Trp53* alone (in *Wap-Cre* control mice) did not cause any significant effect on γH2AX staining in Purkinje cells but led to a substantial reduction in 8-oxo-dG positivity (Figures 6I and 6J), and the low basal level of apoptosis (detected in whole brain sagittal sections) was unaffected (Figure 6K). Co-ablation of *Trp53* with *Palb2* had no effect on any of these three parameters, whereas inactivation of *Trp53* in *Atg7*-CKO and *Palb2*;*Atg7*-CKO mice led to a decrease in all three parameters (Figures 6I-K); however, the results of γH2AX and 8-oxo-dG need to be interpreted with caution because most of the Purkinje cells were lost in mice with wt p53. Taken together, these results demonstrate an important role for p53 in inducing neuronal death upon combined loss of PALB2 and ATG7/autophagy.

### Partial rescue of survival of *Palb2*^Δ/Δ^*;Atg7*^Δ/Δ^ mice by an ROS scavenger

Collectively, results presented above suggest that excessively high oxidative stress, rather than increased DNA damage, in the brain and particularly Purkinje cells underlies the severe neurodegenerative phenotype of *Palb2*;*Atg7*-CKO and *Palb2*^Δ/Δ^*;Atg7*^Δ/Δ^ mice. To test this notion, we treated *Palb2*^Δ/Δ^*;Atg7*^Δ/Δ^ mice with N-acetylcysteine (NAC), an ROS scavenger, and monitored their survival. Compared with untreated mice, NAC-treated animals showed similar survival during the first 35 days or so and then survived significantly longer (Figures 7A and 7B). The divergence suggests that there likely were two different causes of death in the mice and that NAC protects against late death or the second cause, which apparently was neurodegeneration. Note that early death appeared to be caused by tamoxifen toxicity and bacterial infection.

**Figure 7.**
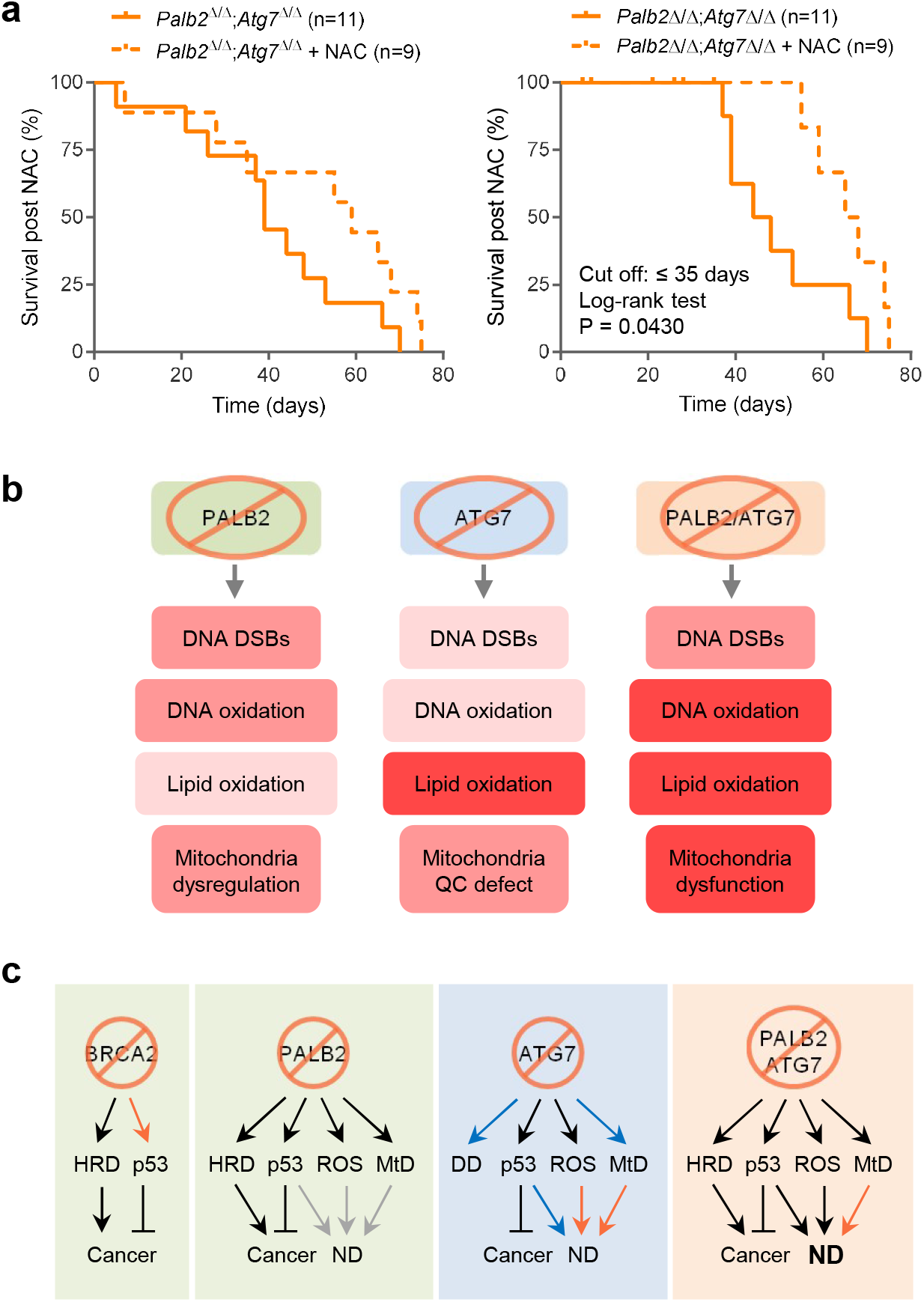
Partial rescue of *Palb2;Atg7*-WBKO mouse survival by an ROS scavenger and proposed events leading to neurodegeneration and cancer in the model mice. (A and B) Survival curves of untreated and NAC-treated *Palb2;Atg7*-WBKO mice. Panel A includes all cause deaths, while in panel B deaths that occurred within the first 35 days were treated as censoring events due to causes unrelated to neurodegeneration. (C) A summary of the impact of PALB2 and ATG7 loss on DNA damage (DSBs), DNA oxidation, lipid oxidation and mitochondria. Colors from light to dark denote mild to strong effects. See texts for details. (D) A proposed model of the mechanisms leading to neurodegeneration and cancer development in the genetically engineered mice used in this study. Black arrows denote strong or well-established effects, blue for moderate effects, and orange and grey for effects that are likely or possible, respectively, but not yet directly proven in the experimental paradigm used. DD, DNA damage; MtD mitochondrial dysfunction.

## Discussion

Autophagy plays important roles in redox homeostasis, mitochondrial quality control, and genome maintenance (Filomeni et al., 2015). It is well established that basal levels of autophagy protect against neurodegeneration, as has been shown by various mouse models with conditional deletion of *Atg7* (Ahmed et al., 2012; Komatsu et al., 2006; Komatsu et al., 2007; Niu et al., 2016). In this study, our results from unintended *Wap-Cre* driven deletion of *Atg7* in the brain and intended *Ubc-Cre-ERT2* driven whole-body knockout of the gene in adult mice both showed that loss of ATG7 in the brain leads to apoptosis of neurons, especially Purkinje cells, motor deficits and shortened survival. Moreover, *Atg7* deletion in the brain led to increased mitochondrial mass, reduced GSH/GSSH ratio, increased DNA and lipid oxidation, as well as increased DNA damage. These results are largely consistent with published reports and further underscore the key roles of autophagy in the above-noted processes and neuronal health.

PALB2 is known to promote HR-mediated DSB repair and antioxidant response (Ma et al., 2012; Xia et al., 2006). In keeping with these known functions, ablation of *Palb2* in the brain led to elevated DNA breaks and DNA oxidation in Purkinje cells (Figures 3A-D). Importantly, our analyses of *Palb2* CKO mouse brains and *PALB2* KO human (DAOY) medulloblastoma cells led to the discovery of a novel role for PALB2 in regulating mitochondrial biogenesis and function. Specifically, loss of PALB2 caused increased mitochondrial mass in both the mouse brain and human DAOY cells but decreased mitochondrial respiration in DAOY cells (Figures 3 and 4). This new discovery adds to the repertoire of PALB2 functions and may have implications in not only neurodegenetation but also tumor suppression. Our studies also establish PALB2 as a suppressor of neurodegeneration in its own right, representing another new function of PALB2.

Interestingly, co-deletion of *Palb2* and *Atg7* in the mouse brain with either *Wap-Cre* or *Ubc-Cre-ERT2* led to a neurodegenerative phenotype that was substantially more severe than that of *Atg7* single deletion mice. When compared with *Atg7*-CKO mice, *Palb2;Atg7*-CKO mice showed significantly accelerated progression of motor dysfunction and imbalance, accompanied by accelerated Purkinje cell loss, and greatly shortened survival (Figure 1). Detailed analyses of mouse brain tissues and human DAOY cells revealed both similarities and differences between the impact of loss of the two proteins on DNA damage, oxidative stress, and mitochondrial homeostasis. Relatively speaking, in the mouse brain, *Palb2* deletion led to larger increases in DSB formation, DNA oxidation, and mitochondrial mass in Purkinje cells, whereas *Atg7* deletion resulted in stronger lipid peroxidation (Figure 3). Human *PALB2*-KO DAOY cells showed larger increases in mitochondrial mass and higher cellular ROS as detected by DCF-DA, while mostly detects H_2_O_2_, whereas *ATG7*-KO cells showed higher levels of mitochondrial superoxide as detected by MitoSox Red (Figure 4). Compared with either single KO cells, *PALB2*;*ATG7*-DKO cells showed even higher cellular ROS, more severe impairment of mitochondrial function, and more spontaneous cell death. These results suggest that PALB2 and ATG7 regulate different aspects of redox and mitochondrial homeostasis and that the severe phenotypes of the double CKO/WBKO mice and the DKO human cells are due to simultaneous loss of complementary functions of ATG7 and PALB2 in antioxidant defense and mitochondrial homeostasis. As neurons, especially Purkinje cells, are sensitive to oxidative stress and energy deprivation, they are also especially vulnerable to combined loss of ATG7/autophagy and PALB2.

Finally, deletion of *Trp53* delayed and alleviated the neurodegenerative phenotype of *Palb2*;*Atg7*-CKO mice and significantly prolonged survival, suggesting that p53 plays a key role in causing this phenotype and reduced survival. Notably, *Trp53* deletion did not affect Purkinje cell number in *Atg7*-CKO mice but restored their number in *Palb2;Atg7*-CKO mice to the same as (but not more than) that in *Atg7*-CKO mice at 6 weeks of age (Figure 6F). This suggests that the loss of Purkinje cells in *Atg7*-CKO mice may not be caused by p53 activation, although p53 activation may lead to apoptosis later or in other neurons, whereas the accelerated Purkinje cell loss in *Palb2*;*Atg7*-CKO mice was largely an *Atg7* null phenotype worsened by further accumulation and/or activation of p53 induced by loss of PALB2.

In summary, our in vivo and cell-based analyses revealed a novel function of PALB2 in regulating mitochondrial homeostasis and further established its less studied role in suppressing ROS, which together distinguish PALB2 from its major binding partner BRCA2 in the DNA damage response and cancer suppression. Moreover, as illustrated in Figures 7C and 7D, we showed that loss of PALB2 and ATG7 lead to different degrees of DNA damage, oxidative stress, mitochondrial abnormality, and p53 induction, all of which may contribute to neurodegeneration, although the impact of any moderate difference in each factor remains to be defined. Still, our results clearly demonstrate that deletion of *Trp53* partially rescues the Purkinje cells in *Palb2;Atg7*-CKO mice and that NAC treatment prolonges survival of *Palb2;Atg7*-WBKO mice. As such, our observations indicate that the severe neurodegenerative phenotypes of *Palb2*;*Atg7* double deletion mice stem to a significant degree from excessive ROS and is at least in part caused by p53-induced neuronal apoptosis. While it is likely that p53 may be induced by DNA damage caused by loss of PALB2’s HR function, whether p53 induction and/or acvtivation is also linked to ROS and mitochondrial dysregulation in the paradigm used awaits further investigation. Additionally, the p53-independent component of the cause of neurodegeneration in mice with *Atg7* deletion also remains to be eclucidated.

## STAR Methods

### Mouse models

The *Palb2^flox2-3^*;*Wap-Cr*e (*Palb2*-CKO) mice were described previously(Huo et al., 2013). They were crossed to strains carrying *Atg7^flox^* (Komatsu et al., 2006), *Brca2^flox11^* (Jonkers et al., 2001), *Trp53^flox2-10^* (Jonkers et al., 2001), or *Ubc-Cre-ERT2* (The Jackson Laboratory) alleles to generate all the genotypes in this study. *Wap-Cre* driven CKO females were all mated to go through two rounds of pregnancy and lactation to induce Cre expression and then monitored for survival and tumor development; males were used for studying neurodegeneration. To generate whole-body somatic *Palb2* and/or *Atg7* knockout in adult mice, 8-10 weeks old males with floxed alleles and *Ubc-Cre-ERT2* were subjected to 5 consecutive daily intraperitoneal injections (unless otherwise specified) of tamoxifen (TAM, 200 μl of 10mg/ml solution per mouse). A PCR based genotyping method was used to validate *Palb2* deletion as described before(Bowman-Colin et al., 2013), and primers used for validation of *Atg7* gene deletion were as follows: forward 5’-aggcagggaggctaaatggt-3’, reverse 5’- gggcgccagttaagaacgat-3’. Diagnostic criteria for neurodegeneration-associated death are 1) a mouse has motor and coordination and balance impairment; 2) loses more than 15% body weight; and 3) no tumors found upon dissection. All animal work were approved by the Institutional Animal Care and Use Committee (IACUC) of the Rutgers Robert Wood Johnson Medical School.

### Motor coordination assessments in mice

Motor coordination was assessed as previously described(Brooks and Dunnett, 2009) with minor modifications. For the footprint test, the forepaws and hindpaws of the mice were coated with red and green nontoxic paints, respectively. Mice were trained to walk along a 50-cm-long, 10-cm-wide, paper-covered runway (with 10-cm-high walls) into an enclosed box. All the mice (5-7 mice per group) were given three runs per day at each time point (6, 9 and 12 weeks of age). A fresh sheet of white paper was placed on the floor of the runway for each run. Footprint patterns were assessed quantitatively by three measurements: stride length, hindbase width and front/hind footprint overlap. Beam balance test was performed using a homebuilt experimental setup of a 50 cm-long, 12 mm-diameter round horizontal beam. At each time point (7, 9, 11, 13, 15, 17, 19 and 20 weeks of age), mice (5-7 per group) were first trained to walk on the beam. For data collection, three trials were performed per animal per day. The time the animal remained on the beam and the steps taken without falling were recorded. Both falling and hanging on the beam were counted as falls (a cushioned pad was used to prevent injury). Mice that successfully walked on the beam for 10 steps without falling were assigned 100 points. For mice that could not pass the “10 steps no fall rule”, we used a walk and stay combined rule in which their score depended on how many steps they could walk and how long they could stay at the spot where they eventually fell off. In this case, number of steps taken and length of stay each accounted for half, or 50 points, of the full score, and each step walked or each 6 seconds of stay was given 5 points; these were then added to yield the score for these mice. The final daily beam walking score was the mean score of the three beam-walking trials.

### Histological Examination and Electron Microscopy

Mice were subjected to cardiac perfusion under deep anethesia with 0.1 M phosphate buffer containing 4% paraformaldehyde. Slides with Meyer’s H&E-stained and immunofluorescent stained midsagittal brain sections were scanned using an Olympus VS120 instrument and images were generated using OlyVIA software for viewing and quantification of Purkinje cells. For electron microscopy, thin sections (70 nm) of the midbrain were prepared and examined by transmission electron microscopy (H7500; Hitachi, Tokyo, Japan). Midbrain tissue sections were analyzed by quantifying at least 5 images at 3800×magnification using ImageJ to measure mitochondrial mass. Two mice of each genotype were analyzed. Mitochondrial intensity index (MDI) was calculated as follows: MDI (mitochondria density index) = na/A, where A = area of each image, a = area of each mitochondria, and n = number of mitochondria in the image.

### Immunohistochemistry (IHC)

Tissues were fixed in 10% buffered formalin solution for 24-72 hrs and then transferred to 70% ethanol prior to further processing. Paraffin-embedded block production and sectioning were conducted by The Histopathology Shared Resources of Rutgers Cancer Institute of New Jersey. IHC was performed on 5 μm sections as described before(Simhadri et al., 2014). Paraffin sections were stained with antibodies against p53 (CM5, 1:500; Leica Biosystems), γ-H2AX (1:300; Millipore, 05-636), 8-oxo-dG (1:2000; Trevigen); MT-CO1 (1:300, Abcam, ab14705) and Mn-SOD2 (1:200, EMD Millipore, 06-984). Brain tissue from three mice in each group were analyzed by quantifying Purkinje cells of whole brain sagittal sections at 200× magnification. For γ-H2AX staining, ≥1 foci/cell were counted as positive; for 8-oxo-dG, both light and dark brown staining were counted as positive; MT-CO1 staining was scored as 1, 2 and 3 based on staining intensity. At least 100 Purkinje cells or all Purkinje cells in a brain section (when the total number in the section was less than 100) were counted.

### GSH/GSSG measurement

GSH and GSSG contents in the brain were measured as part of a comprehensive metabolomics analysis using liquid chromatography-mass spectrometry (LC-MS). In brief, brain tissues were collected from the *Palb2*, *Atg7*, and *Palb2;Atg7* WBKO mice at 4 weeks post tamoxifen treatment and snap frozen in liquid N2. Frozen tissue samples were weighed (~20-30 mg) and ground in liquid N2 using Cryomill (Retsh). The powdered samples were then mixed with methanol: acetonitrile: water (40:40:20) solution containing 0.5% formic acid, followed by 10 sec vortexing, 10 min incubation on ice and centrifugation at 16,000 × g at 4°C. The supernatents were collected, neutralized with 50μL/mL of 15% ammonium bicarbonate, and then centrifuged at 16,000 × g for 10 min. The final supernatants were transferred to LC–MS autosampler vials and sent for LC-MS analysis which was done using Q Exactive Plus hybrid quadrupole orbitrap mass spectrometer (Thermo Fisher scientific). Raw data analysis was done using *Maven* v.707.

### Treatment of mice with N-Acetyl-Cysteine (NAC)

Eight to ten weeks old *Ubc-Cre-ERT2* model mice were subjected to 4 daily injections of tamoxifen to delete genes of interest and then separated into 2 groups. One group was treated with sterile filtered water supplemented with 1% NAC (Sigma A9165) (pH ~7.4) and the other group was given control water. The water was changed every 2-3 days and the mice were monitored for survival.

### Western blotting

Cells were lysed with NETNG-400 (400mM NaCl, 1 mM EDTA, 20 mM Tris-HCl [pH7.5], 0.5% Non-Idet P-40, and 10% glycerol) with Complete® protease inhibitor cocktail (Roche). Tissues were homogenized in the same buffer by a TissueRuptor (Qiagen) followed by sonication. Samples were resolved on 4-12% or 4-20%Tris-glycine SDS-polyacrylamide gels and transferred to a nitrocellulose membrane (0.45 micron, Bio-Rad) overnight at 4°C. Resolved proteins were detected following standard procedures using the following antibodies: ATG7 (1:2000; Sigma A2856), LC3 (1:1500; Novus Biologicals NB600-1384), p62 (1:10000; Abcam, ab109012), PALB2 M11 (1:2000, homemade), Mn-SOD2 (1:2000, EMD Millipore, 06-984), MT-CO1 (1:2000, Abcam, ab14705), MT-CO2 (1:1000, Abcam, ab198286), GAPDH (1:1000; Santa Cruz Biotechnology sc-365062), and β-Actin (1:5000; Sigma A1978). Blots were developed using Immobilon Western Chemiluminescent HRP Substrate (EMD Millipore).

### Terminal deoxynucleotidyltransferase dUTP nick end labeling (TUNEL) assay

Staining was performed on 5 μm sections using the DeadEnd™ Fluorometric TUNEL System (Promega) according to the manufacturer’s instructions, and all apoptotic cells in the cerebellum of each midsagittal section were counted.

### CRISPR/Cas9 knock out of *PALB2* and *ATG7*

Human DAOY medulloblastoma cells were purchased from ATCC and cultured at 37°C in DMEM/F12 (#8437, Sigma) supplemented with 10% FBS and 1% penicillin/streptomycin, in a humidified incubator with 5% CO2. pSpCas9-2A-GFP (PD1301) V2.0 constructs containing gRNAs targeting *PALB2* and pSpCas9(BB)-2A-Puro (PX459) V2.0 constructs containing gRNAs targeting *ATG7* were transfected into DAOY cells using X-tremeGENE 9 (Roche, Mannheim, Germany). Single cell clones were isolated by puromycin selection or fluorescence activated cell sorting (FACS). Clones were first assayed by western blotting for complete loss of protein expression, and genomic DNA of candidate clones were then sequenced to verify the disruption of the genes. The target gRNA sequences against PALB2 are: 5’-gccttcaggtaagtgaatcg-3’ and 5’-ctagcgtgcccaaagagctg-3’, against ATG7 is 5’-gctgccagctcgcttaacat-3’.

### Measurements of ROS, mitochondrial mass and mitochondrial membrane potential

Cells were washed with PBS and then incubated at dark for 30 min with phenol red–free DMEM with 10% FBS and 100 μmol/L 2’,7’-dichlorofluorescein diacetate (DCF-DA; D6883, Sigma) or 2.5 μmol/L MitoSOX™ Red (M36008, Invitrogen) for ROS measurement, 200 nmol/L of MitoTracker® Green FM (M7512, Invitrogen) for mitochondrial mass measurement and 200 nmol/L of MitoTracker® Red CMXRos (M7512, Invitrogen) for mitochondrial membrane potential measurement. After incubation, cells were trypsinized, spun down, and resuspended in PBS at a density of approximately 2×10^5^ cells per mL. Signals were analyzed by flow cytometry.

### Assessment of oxygen consumption rate (OCR)

OCR of control and KO DAOY cells were measured using a Seahorse Biosciences extracellular flux analyzer (XF24). Cells were seeded at 2.0 × 10^4^ cells per well in the XF24 plates overnight prior to XF assay. Real-time OCR measurements were performed in DMEM/F12 with 2 mM glutamine, or 1 mM dimethylα-KG for 3 h, and measurements were taken every 15 min. Relative OCR (percentage) was normalized to the 0-min time point.

### Statistical analyses

Comparisons of survival curves were made using the log-rank test(Crowley and Breslow, 1984). Unpaired two-tailed t-test was used for column analysis with GraphPad Prism 8.4.3. Significance is denoted as follows: ns, p>0.05; *, p≤0.05; **, p≤0.01. ***, p≤0.001. All error bars represent standard errors (SE).

## Acknowledgements

This work was supported by the National Cancer Institute R01CA188096 (Xia and White) and R01CA138804 (Xia). This research was also supported by the Biospecimen Repository and Histopathology Service Shared Resource, the Flow Cytometry and Cell Sorting Shared Resource, and the Metabolomics Shared Resource of Rutgers Cancer Institute of New Jersey, which are supported in part by P30CA072720 from the National Cancer Institute. M.M.M. is the William Dow Lovett Professor of Neurology and is supported by NIH grants NS101134, NS096032, and NS11692, the Michael J. Fox Foundation for Parkinson’s Research and the American Parkinson Disease Association.

## References

Ahmed, I., Liang, Y., Schools, S., Dawson, V.L., Dawson, T.M., and Savitt, J.M. (2012). Development and characterization of a new Parkinson’s disease model resulting from impaired autophagy. J Neurosci 32, 16503–16509.

Aquilano, K., Baldelli, S., and Ciriolo, M.R. (2014). Glutathione: new roles in redox signaling for an old antioxidant. Front Pharmacol 5, 196.

Ashrafi, G., and Schwarz, T.L. (2013). The pathways of mitophagy for quality control and clearance of mitochondria. Cell Death Differ 20, 31–42.

Baxter, P.S., and Hardingham, G.E. (2016). Adaptive regulation of the brain’s antioxidant defences by neurons and astrocytes. Free Radic Biol Med 100, 147–152.

Bjorkoy, G., Lamark, T., Brech, A., Outzen, H., Perander, M., Overvatn, A., Stenmark, H., and Johansen, T. (2005). p62/SQSTM1 forms protein aggregates degraded by autophagy and has a protective effect on huntingtin-induced cell death. Journal of Cell Biology 171, 603–614.

Bowman-Colin, C., Xia, B., Bunting, S., Klijn, C., Drost, R., Bouwman, P., Fineman, L., Chen, X.X., Culhane, A.C., Cai, H., et al. (2013). Palb2 synergizes with Trp53 to suppress mammary tumor formation in a model of inherited breast cancer. P Natl Acad Sci USA 110, 8632–8637.

Boya, P., Codogno, P., and Rodriguez-Muela, N. (2018). Autophagy in stem cells: repair, remodelling and metabolic reprogramming. Development 145.

Brooks, S.P., and Dunnett, S.B. (2009). Tests to assess motor phenotype in mice: a user’s guide. Nat Rev Neurosci 10, 519–529.

Butterfield, D.A., Reed, T., Newman, S.F., and Sultana, R. (2007). Roles of amyloid beta-peptide-associated oxidative stress and brain protein modifications in the pathogenesis of Alzheimer’s disease and mild cognitive impairment. Free Radical Bio Med 43, 658–677.

Canugovi, C., Misiak, M., Ferrarelli, L.K., Croteau, D.L., and Bohr, V.A. (2013). The role of DNA repair in brain related disease pathology. DNA Repair 12, 578–587.

Collaborators, G.B.D.N. (2019). Global, regional, and national burden of neurological disorders, 1990-2016: a systematic analysis for the Global Burden of Disease Study 2016. Lancet Neurol 18, 459–480.

Crowley, J., and Breslow, N. (1984). Statistical analysis of survival data. Annu Rev Public Health 5, 385–411.

Filomeni, G., De Zio, D., and Cecconi, F. (2015). Oxidative stress and autophagy: the clash between damage and metabolic needs. Cell Death Differ 22, 377–388.

Forrester, S.J., Kikuchi, D.S., Hernandes, M.S., Xu, Q., and Griendling, K.K. (2018). Reactive Oxygen Species in Metabolic and Inflammatory Signaling. Circ Res 122, 877–902.

Friedman, L.G., Lachenmayer, M.L., Wang, J., He, L., Poulose, S.M., Komatsu, M., Holstein, G.R., and Yue, Z. (2012). Disrupted autophagy leads to dopaminergic axon and dendrite degeneration and promotes presynaptic accumulation of alpha-synuclein and LRRK2 in the brain. J Neurosci 32, 7585–7593.

Guo, J.Y., Teng, X., Laddha, S.V., Ma, S., Van Nostrand, S.C., Yang, Y., Khor, S., Chan, C.S., Rabinowitz, J.D., and White, E. (2016). Autophagy provides metabolic substrates to maintain energy charge and nucleotide pools in Ras-driven lung cancer cells. Genes Dev 30, 1704–1717.

Guo, J.Y., and White, E. (2013). Autophagy is required for mitochondrial function, lipid metabolism, growth, and fate of KRAS(G12D)-driven lung tumors. Autophagy 9, 1636–1638.

Guo, J.Y., Xia, B., and White, E. (2013). Autophagy-mediated tumor promotion. Cell 155, 1216–1219.

Hara, T., Nakamura, K., Matsui, M., Yamamoto, A., Nakahara, Y., Suzuki-Migishima, R., Yokoyama, M., Mishima, K., Saito, I., Okano, H., et al. (2006). Suppression of basal autophagy in neural cells causes neurodegenerative disease in mice. Nature 441, 885–889.

Huo, Y., Cai, H., Teplova, I., Bowman-Colin, C., Chen, G., Price, S., Barnard, N., Ganesan, S., Karantza, V., White, E., et al. (2013). Autophagy opposes p53-mediated tumor barrier to facilitate tumorigenesis in a model of PALB2-associated hereditary breast cancer. Cancer discovery 3, 894–907.

Inoue, K., Rispoli, J., Kaphzan, H., Klann, E., Chen, E.I., Kim, J., Komatsu, M., and Abeliovich, A. (2012). Macroautophagy deficiency mediates age-dependent neurodegeneration through a phospho-tau pathway. Mol Neurodegener 7, 48.

Jacobsen, P.F., Jenkyn, D.J., and Papadimitriou, J.M. (1985). Establishment of a human medulloblastoma cell line and its heterotransplantation into nude mice. Journal of neuropathology and experimental neurology 44, 472–485.

Jonkers, J., Meuwissen, R., van der Gulden, H., Peterse, H., van der Valk, M., and Berns, A. (2001). Synergistic tumor suppressor activity of BRCA2 and p53 in a conditional mouse model for breast cancer. Nat Genet 29, 418–425.

Karsli-Uzunbas, G., Guo, J.Y., Price, S., Teng, X., Laddha, S.V., Khor, S., Kalaany, N.Y., Jacks, T., Chan, C.S., Rabinowitz, J.D., et al. (2014). Autophagy is required for glucose homeostasis and lung tumor maintenance. Cancer discovery 4, 914–927.

Kimmelman, A.C., and White, E. (2017). Autophagy and Tumor Metabolism. Cell Metab 25, 1037–1043.

Klionsky, D.J., and Schulman, B.A. (2014). Dynamic regulation of macroautophagy by distinctive ubiquitin-like proteins. Nat Struct Mol Biol 21, 336–345.

Kokoszka, J.E., Coskun, P., Esposito, L.A., and Wallace, D.C. (2001). Increased mitochondrial oxidative stress in the Sod2 (+/−) mouse results in the age-related decline of mitochondrial function culminating in increased apoptosis. Proc Natl Acad Sci U S A 98, 2278–2283.

Komatsu, M., Waguri, S., Chiba, T., Murata, S., Iwata, J., Tanida, I., Ueno, T., Koike, M., Uchiyama, Y., Kominami, E., et al. (2006). Loss of autophagy in the central nervous system causes neurodegeneration in mice. Nature 441, 880–884.

Komatsu, M., Wang, Q.J., Holstein, G.R., Friedrich, V.L., Jr., Iwata, J., Kominami, E., Chait, B.T., Tanaka, K., and Yue, Z. (2007). Essential role for autophagy protein Atg7 in the maintenance of axonal homeostasis and the prevention of axonal degeneration. Proc Natl Acad Sci U S A 104, 14489–14494.

Li, L., Zhang, X., and Le, W. (2008). Altered macroautophagy in the spinal cord of SOD1 mutant mice. Autophagy 4, 290–293.

Ma, J., Cai, H., Wu, T., Sobhian, B., Huo, Y., Alcivar, A., Mehta, M., Cheung, K.L., Ganesan, S., Kong, A.N., et al. (2012). PALB2 interacts with KEAP1 to promote NRF2 nuclear accumulation and function. Molecular and cellular biology 32, 1506–1517.

Murphy, M.P. (2009). How mitochondria produce reactive oxygen species. Biochem J 417, 1–13.

Nahapetyan, H., Moulis, M., Grousset, E., Faccini, J., Grazide, M.H., Mucher, E., Elbaz, M., Martinet, W., and Vindis, C. (2019). Altered mitochondrial quality control in Atg7-deficient VSMCs promotes enhanced apoptosis and is linked to unstable atherosclerotic plaque phenotype. Cell Death Dis 10, 119.

Niu, X.Y., Huang, H.J., Zhang, J.B., Zhang, C., Chen, W.G., Sun, C.Y., Ding, Y.Q., and Liao, M. (2016). Deletion of autophagy-related gene 7 in dopaminergic neurons prevents their loss induced by MPTP. Neuroscience 339, 22–31.

Nixon, R.A., Wegiel, J., Kumar, A., Yu, W.H., Peterhoff, C., Cataldo, A., and Cuervo, A.M. (2005). Extensive involvement of autophagy in Alzheimer disease: an immuno-electron microscopy study. Journal of neuropathology and experimental neurology 64, 113–122.

Poillet-Perez, L., and White, E. (2019). Role of tumor and host autophagy in cancer metabolism. Genes Dev 33, 610–619.

Rak, M., Benit, P., Chretien, D., Bouchereau, J., Schiff, M., El-Khoury, R., Tzagoloff, A., and Rustin, P. (2016). Mitochondrial cytochrome c oxidase deficiency. Clin Sci (Lond) 130, 393–407.

Ravikumar, B., Vacher, C., Berger, Z., Davies, J.E., Luo, S.Q., Oroz, L.G., Scaravilli, F., Easton, D.F., Duden, R., O’Kane, C.J., et al. (2004). Inhibition of mTOR induces autophagy and reduces toxicity of polyglutamine expansions in fly and mouse models of Huntington disease. Nat Genet 36, 585–595.

Ruzankina, Y., Pinzon-Guzman, C., Asare, A., Ong, T., Pontano, L., Cotsarelis, G., Zediak, V.P., Velez, M., Bhandoola, A., and Brown, E.J. (2007). Deletion of the developmentally essential gene ATR in adult mice leads to age-related phenotypes and stem cell loss. Cell Stem Cell 1, 113–126.

Simhadri, S., Peterson, S., Patel, D.S., Huo, Y., Cai, H., Bowman-Colin, C., Miller, S., Ludwig, T., Ganesan, S., Bhaumik, M., et al. (2014). Male fertility defect associated with disrupted BRCA1-PALB2 interaction in mice. J Biol Chem 289, 24617–24629.

Singh, A., Kukreti, R., Saso, L., and Kukreti, S. (2019). Oxidative Stress: A Key Modulator in Neurodegenerative Diseases. Molecules 24.

Son, J.H., Shim, J.H., Kim, K.H., Ha, J.Y., and Han, J.Y. (2012). Neuronal autophagy and neurodegenerative diseases. Experimental & molecular medicine 44, 89–98.

Strohecker, A.M., Guo, J.Y., Karsli-Uzunbas, G., Price, S.M., Chen, G.J., Mathew, R., McMahon, M., and White, E. (2013). Autophagy sustains mitochondrial glutamine metabolism and growth of BrafV600E-driven lung tumors. Cancer discovery 3, 1272–1285.

Trentesaux, C., Fraudeau, M., Pitasi, C.L., Lemarchand, J., Jacques, S., Duche, A., Letourneur, F., Naser, E., Bailly, K., Schmitt, A., et al. (2020). Essential role for autophagy protein ATG7 in the maintenance of intestinal stem cell integrity. Proc Natl Acad Sci U S A 117, 11136–11146.

Vogiatzi, T., Xilouri, M., Vekrellis, K., and Stefanis, L. (2008). Wild type alpha-synuclein is degraded by chaperone-mediated autophagy and macroautophagy in neuronal cells. Journal of Biological Chemistry 283, 23542–23556.

Wagner, K.U., Wall, R.J., St-Onge, L., Gruss, P., Wynshaw-Boris, A., Garrett, L., Li, M., Furth, P.A., and Hennighausen, L. (1997). Cre-mediated gene deletion in the mammary gland. Nucleic Acids Res 25, 4323–4330.

Wang, C., Chen, S., Yeo, S., Karsli-Uzunbas, G., White, E., Mizushima, N., Virgin, H.W., and Guan, J.L. (2016). Elevated p62/SQSTM1 determines the fate of autophagy-deficient neural stem cells by increasing superoxide. J Cell Biol 212, 545–560.

Watts, M.E., Pocock, R., and Claudianos, C. (2018). Brain Energy and Oxygen Metabolism: Emerging Role in Normal Function and Disease. Front Mol Neurosci 11, 216.

Wei, H., Wei, S., Gan, B., Peng, X., Zou, W., and Guan, J.L. (2011). Suppression of autophagy by FIP200 deletion inhibits mammary tumorigenesis. Genes Dev 25, 1510–1527.

Xia, B., Sheng, Q., Nakanishi, K., Ohashi, A., Wu, J., Christ, N., Liu, X., Jasin, M., Couch, F.J., and Livingston, D.M. (2006). Control of BRCA2 cellular and clinical functions by a nuclear partner, PALB2. Mol Cell 22, 719–729.

Yang, W.S., SriRamaratnam, R., Welsch, M.E., Shimada, K., Skouta, R., Viswanathan, V.S., Cheah, J.H., Clemons, P.A., Shamji, A.F., Clish, C.B., et al. (2014). Regulation of ferroptotic cancer cell death by GPX4. Cell 156, 317–331.

Yang, Y., Karsli-Uzunbas, G., Poillet-Perez, L., Sawant, A., Hu, Z.S., Zhao, Y., Moore, D., Hu, W., and White, E. (2020). Autophagy promotes mammalian survival by suppressing oxidative stress and p53. Genes Dev 34, 688–700.

Yoshida, R. (2020). Hereditary breast and ovarian cancer (HBOC): review of its molecular characteristics, screening, treatment, and prognosis. Breast Cancer.

Zhang, F., Ma, J.L., Wu, J.X., Ye, L., Cai, H., Xia, B., and Yu, X.C. (2009). PALB2 Links BRCA1 and BRCA2 in the DNA-Damage Response. Curr Biol 19, 524–529.

Zong, W.X., Rabinowitz, J.D., and White, E. (2016). Mitochondria and Cancer. Mol Cell 61, 667–676.

Zorov, D.B., Juhaszova, M., and Sollott, S.J. (2014). Mitochondrial reactive oxygen species (ROS) and ROS-induced ROS release. Physiol Rev 94, 909–950.

