## Supplementary Figures 1 and 2 for "PALB2 maintains redox and mitochondrial homeostasis in the brain and cooperates with ATG7 to suppress p53 dependent neurodegeneration"

Huo et al., Supplemental Figure 1

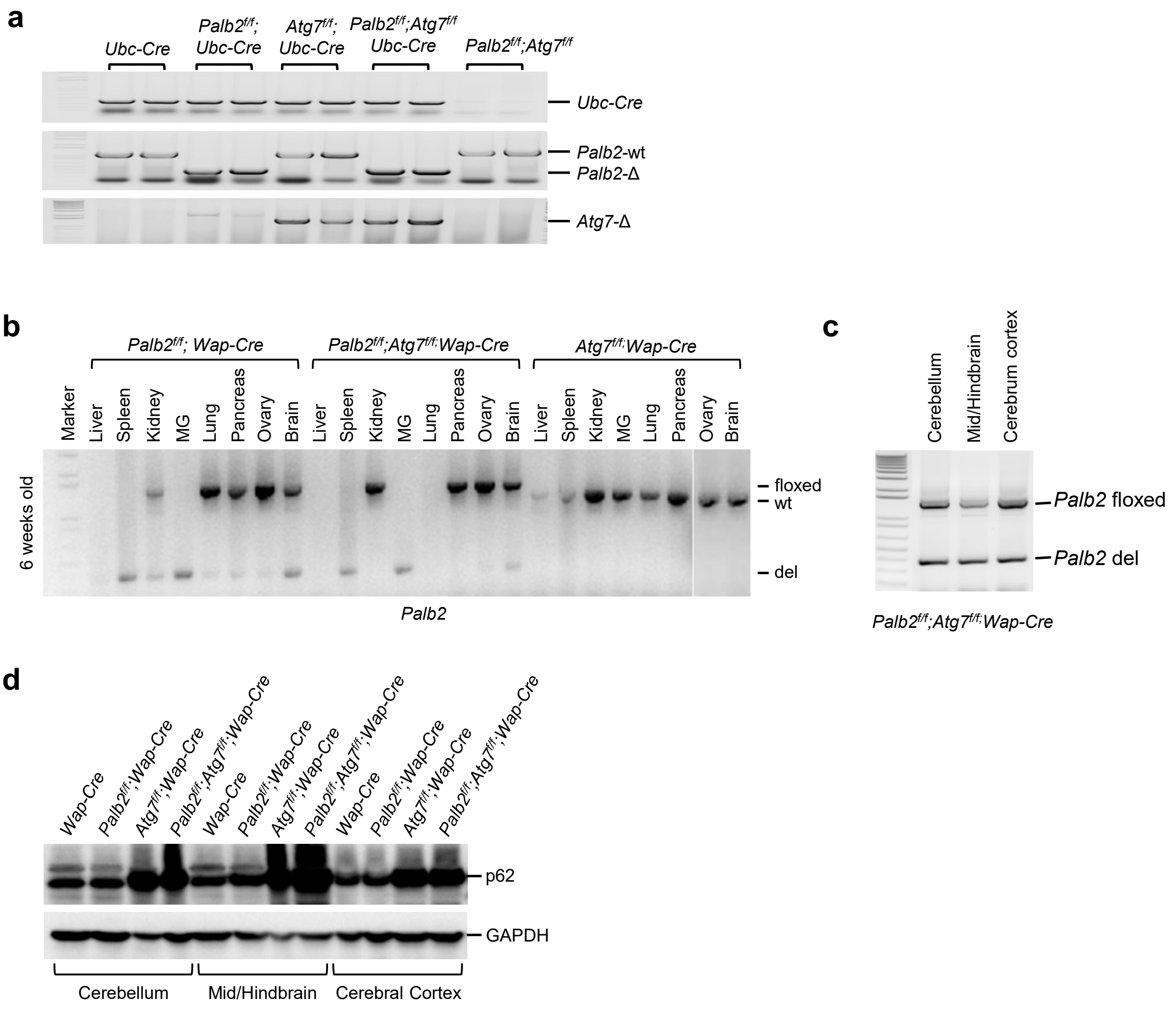


**Figure S1. Detection of *Palb2* and/or *Atg7* deletion in *Wap-cre* or *Ubc-Cre-ERT2* model mice. a** PCR detection of tamoxifen induced, Ubc-Cre-ERT2-mediated *Palb2* or *Atg7* deletion in the whole brain. **b** PCR detection of *Palb2* deletion in different tissues of 6 weeks *Palb2*$^{f/f}$*;Wap-cre* , *Atg7*$^{f/f}$*;Wap-cre* and *Palb2*$^{f/f}$*;Atg7*$^{f/f}$*;Wap-cre* mice. **c** PCR detection of *Palb2* deletion in brain regions of *Palb2*$^{f/f}$*;Atg7*$^{f/f}$*;Wap-cre* mice. **d** Representative western blots showing p62 accumulation in the cerebellum, mid/hindbrain and cerebrum cortex of 6 weeks old *Wap-cre*, *Palb2*$^{f/f}$*;Wap-cre, Atg7*$^{f/f}$*;Wap-cre* and *Palb2*$^{f/f}$*;Atg7*$^{f/f}$*;Wap-cre* mice. GAPDH was used as a loading control.

Huo et al., Supplemental Figure 2

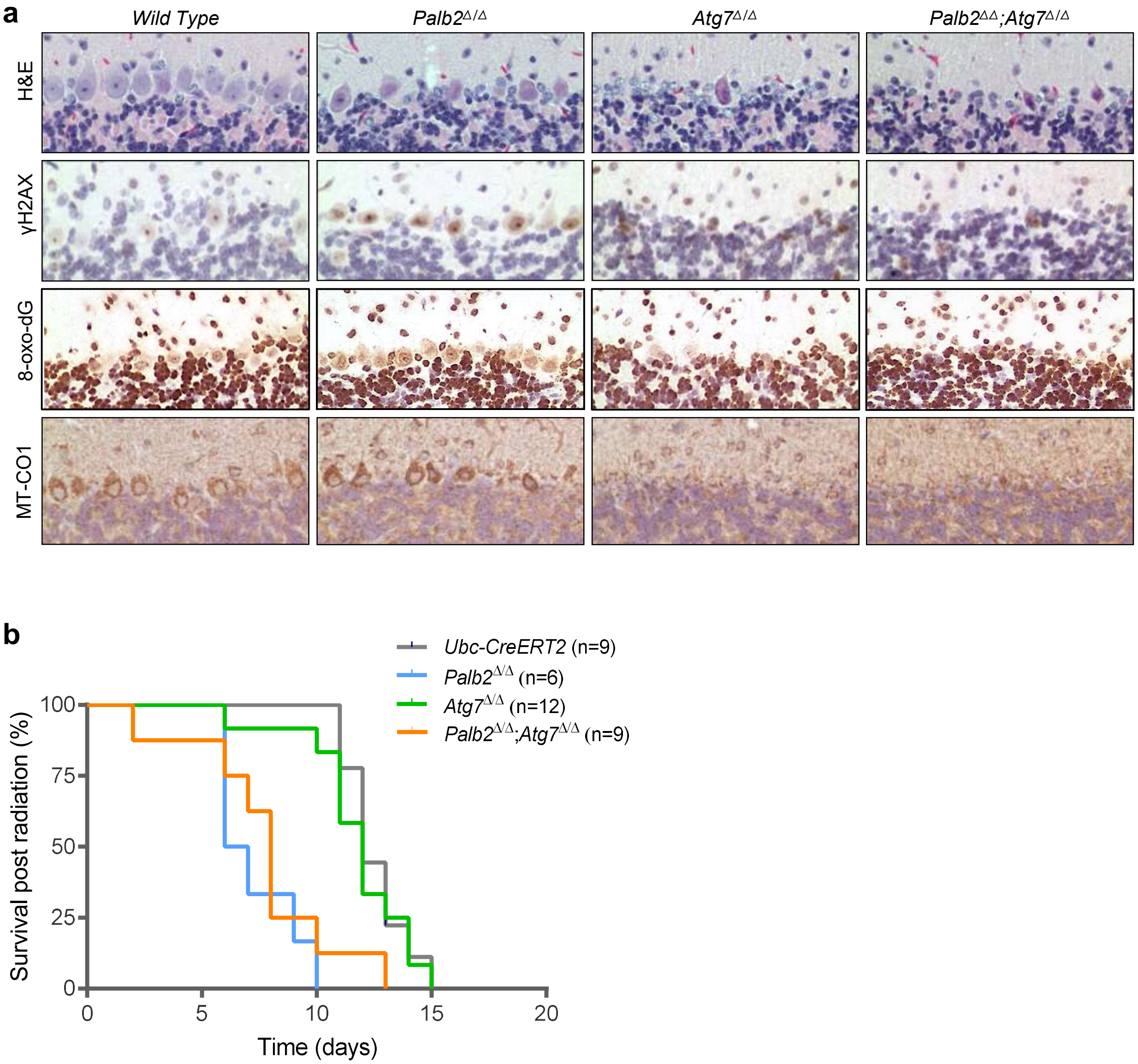


**Figure S2. Markers of DNA damage, oxidative stress and mitochondria in *Palb2*, *Atg7* and *Palb2;Atg7*-WBKO mice and the sensitivity of the mice to radiation. a** Representative H&E staining and IHC staining of γH2AX, 8-oxo-dG MT-CO1 in Purkinje cells and surrounding cells of *UBC-Cre-ERT2*, *Palb2*$^{\Delta\Delta}$, *Atg7*$^{\Delta\Delta}$ *and Palb2*$^{\Delta\Delta}$;*Atg7*$^{\Delta\Delta}$ mice. **b** Survival curves of the above mice after 10 Gy of whole-body γ-radiation.
